# Gbx2 controls amacrine cell dendrite stratification through Robo1/2 receptors

**DOI:** 10.1101/2023.08.03.551861

**Authors:** Patrick C. Kerstein, Yessica Santana Agreda, Bridget M. Curran, Le Ma, Kevin M. Wright

## Abstract

Within the neuronal classes of the retina, amacrine cells (ACs) exhibit the greatest neuronal diversity in morphology and function. We show that the selective expression of the transcription factor *Gbx2* is required for cell fate specification and dendritic stratification of an individual AC subtype in the mouse retina. We identify Robo1 and Robo2 as downstream effectors that when deleted, phenocopy the dendritic misprojections seen in *Gbx2* mutants. Slit1 and Slit2, the ligands of Robo receptors, are localized to the OFF layers of the inner plexiform layer where we observe the dendritic misprojections in both *Gbx2* and *Robo1/2* mutants. We show that Robo receptors also are required for the proper dendritic stratification of additional AC subtypes, such as Vglut3+ ACs. These results show both that Gbx2 functions as a terminal selector in a single AC subtype and identify Slit-Robo signaling as a developmental mechanism for ON-OFF pathway segregation in the retina.

## INTRODUCTION

Neural circuit assembly requires the coordination of genetic pathways that execute multiple tasks: differentiation of the correct cell types, formation of axon and dendrite morphology, and patterning of synaptic connections. Disruption of these developmental processes, even at the level of a single neuronal cell type, can lead to dysfunction in the nervous system (Kulkarni and Firestein, 2012; Lefebvre et al., 2015; Morrie and Feller, 2016). Terminal selector transcription factors guide the proper development of the nervous system by regulating the expression of effector genes necessary for axon guidance and dendrite development, synaptogenesis, neurotransmitter synthesis, and neuronal function (Holguera and Desplan, 2018; Peng, 2023). Identifying which transcription factors control the development of specific neuronal cell types and what genes they regulate is important for both understanding the fundamentals of neural development and the mechanisms necessary for tissue regeneration and repair.

In the retina, recent work has identified several of the transcription factors that control retinal ganglion cell (RGC) development. *Tbr1*, *Tbr2*, and *Satb2* all control effector genes necessary for the cell fate, morphology, and connectivity of specific sets of RGC subtypes (Kiyama et al., 2019; Liu et al., 2018; Peng, 2023; Peng et al., 2020; Sweeney et al., 2014). The genetic mechanisms are likely even more complex for retinal neuron classes that exhibit greater diversity such as amacrine cells (ACs), which consist of 63 molecularly distinct subtypes (Yan et al., 2020). ACs are inhibitory interneurons in the inner retina that aid in encoding specific features of visual stimuli. Each AC subtype has a unique dendritic morphology that is closely tied to its function (Diamond, 2017; Morrie and Feller, 2016). Disruption of dendrite development of even a single AC subtype can lead to a change in visual circuit function. For example, in starburst amacrine cells (SACs), deletion of the transcription factor *Sox2* causes migration and dendrite stratification errors (Whitney et al., 2014) and disrupts the physiological function of the cell (Stincic et al., 2018). Deletion of cell-surface proteins that are critical for SAC dendrite morphology and stratification leads to abnormal SAC function (Kostadinov and Sanes, 2015; Lefebvre et al., 2012; Morrie and Feller, 2018; Soto et al., 2019; Sun et al., 2013). In other AC subtypes, transcription factors such as *NeuroD6*, *Lhx9*, *Prdm13*, *Isl1*, and *Fezf1* regulate aspects of differentiation, cell migration, dendrite stratification, and neurotransmitter expression (Balasubramanian et al., 2018; Elshatory et al., 2007; Goodson et al., 2018; Kay et al., 2011; Peng et al., 2020). However, the genetic basis of cell fate specification and dendrite development has only been explored in a few of the 63 AC subtypes. Here, we focus on the development of the recently identified Gbx2+ AC and identify genes important for regulating its cell fate and dendrite stratification.

Gbx2 is a homeobox transcription factor that controls neural patterning, cell migration, and axon guidance in the developing central nervous system (Chatterjee et al., 2012; Chen et al., 2010; Luu et al., 2011; Mallika et al., 2015). In the retina, a *Gbx2^CreERT2^* knock-in mouse line labels two molecularly, morphologically, and functionally distinct AC subtypes that stratify their dendrites in either sublamina 3 (S3) or sublamina 5 (S5) of the inner plexiform layer (IPL) (Kerstein et al., 2020). However, only the S3-stratifying Gbx2+ AC subtype expresses *Gbx2* at an appreciable level (Kerstein et al., 2020; Macosko et al., 2015; Yan et al., 2020). S3-Gbx2+ ACs have a unique dorsotemporal-oriented asymmetric dendritic morphology and a clear stratification pattern making it an ideal model for studying the genetic basis of cell fate and dendrite development.

In this study, we show that *Gbx2* is required for the initial specification and subsequent dendrite stratification of S3-Gbx2+ ACs. Further, we identify Robo1 and Robo2 receptors as Gbx2 effector genes that regulate dendrite stratification. In both *Gbx2* and *Robo1/2* mutant retinas, Gbx2+ ACs send ectopic dendrites to the upper (OFF) layers of the IPL, where OFF bipolar cells expressing the repulsive Slit1 and Slit2 ligands stratify their axon terminals. Robo signaling is also necessary for dendrite targeting in other ACs, as the S3-stratifying Vglut3+ AC exhibit similar ectopic dendritic projections in the upper layers of the IPL. Together, these results identify a molecular pathway that regulates AC subtype specification and dendrite lamination and identify a role for Robo receptors in mediating dendrite development by preventing ON layer dendrites from targeting the OFF layers of the IPL in the mammalian retina.

## RESULTS

### The dendrites of S3-Gbx2+ ACs develop in two stages: laminar stratification and dendritic arbor elaboration

We previously identified a subtype of non-GABAergic, non-Glycinergic Gbx2+ amacrine cells (ACs) with dorsotemporal-oriented asymmetric dendrites that stratify in sublamina 3 (S3) of the inner plexiform layer (IPL) (Kerstein et al., 2020). To better understand the development of S3-Gbx2+ ACs we defined when and where their dendrites develop within the IPL of the retina. EdU birthdating showed that S3-Gbx2+ ACs are born between E12 and E18, with the peak of their genesis occurring between E14-E16 (Fig. S1). To capture the full morphological development of S3-Gbx2+ ACs, we analyzed retinal cross-sections from *Gbx2^CreERT2-IRES-EGFP^* mice at several developmental timepoints: P1, P4, P7, and P14 (Chen et al. 2009; Kerstein et al. 2020) (Fig. 1a-h) (Chen et al., 2009; Kerstein et al., 2020). At P1, EGFP+ S3-Gbx2+ ACs have begun to extend their dendrites into the IPL, however their arbors lack clear stratification (Fig. 1a, e). At this timepoint, the somas of the S3-Gbx2+ ACs are still in the process of splitting between the inner nuclear layer (INL) and ganglion cell layer (GCL); this lags behind the previous reported development of SACs by about 1 day (Ray et al. 2018). At P4, the dendrites of the Gbx2+ ACs are restricted to the lower layers of the IPL (Fig 1b, f), and by P7 the dendrites exhibit a clear stratification within the center of the IPL (Fig 1c, g). At P14, when the IPL has expanded in size, the Gbx2+ AC dendrites remain stratified in the central part of the IPL (Fig. 1d, h). These data suggest that Gbx2+ ACs stratify their dendrites in the correct layer of the IPL between P4 and P7.

**Figure 1.**
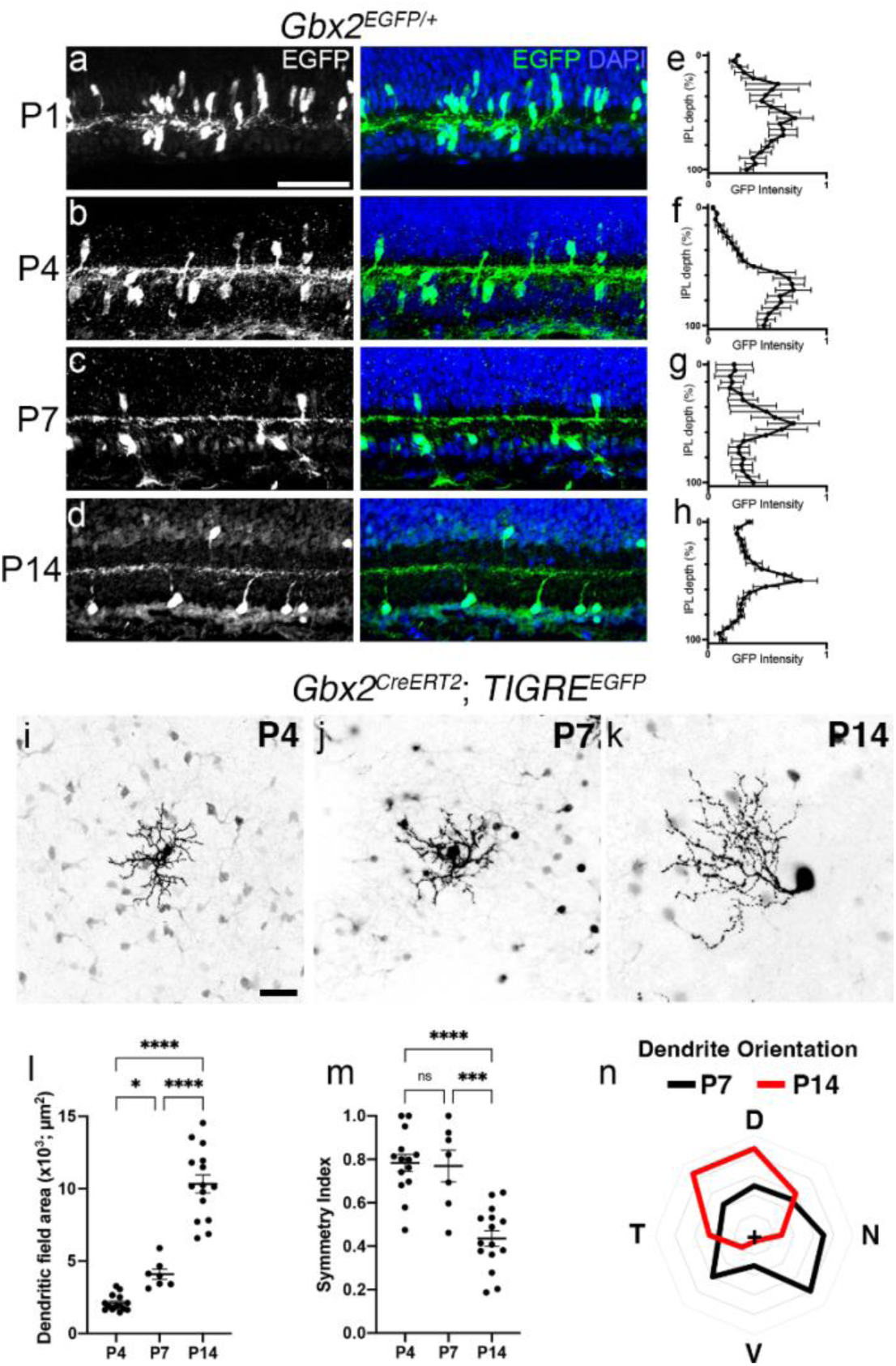
The dendrites of Gbx2+ amacrine cells (ACs) develop over the first two postnatal weeks. (**a-d**) Retinal cross-sections at multiple developmental timepoints from the *Gbx2^EGFP^* mouse line. Left, EGFP-labeling of Gbx2+ ACs and their dendrites at (**a**) P1, (**b**) P4, (**c**) P7, and (**d**) P14. Middle, merged image with DAPI to highlight cellular and neuropil layers of the retina. (e-h) Fluorescence intensity plots through the depth of the IPL showing dendrite localization from P1-P14. Dendrite morphology of S3-Gbx2+ ACs *en face* at (**i**) P4, (**j**) P7, and (**k**) P14 timepoints in retinal flat mounts. Gbx2+ ACs are sparsely labeled using the *Gbx2^CreERT2^, TIGRE^LSL-EGFP^* (Ai140D) mouse line and low dose tamoxifen (0.5 µg) at P0-P1. Quantification of Gbx2+ AC dendritic arbor (**l**) area and (**m**) symmetry at several developmental timepoints. (**n**) Dendrites of Gbx2+ ACs do not show a specific orientation at P7 (black) but orient in the dorsal-temporal direction by P14 (red). ns, not significant, *p<0.05, ***p<0.001, and ****p<0.0001 using an ordinary One-Way ANOVA with Multiple Comparisons Test. Scale bars (**a, i**), 50μm.

Next, we determined when S3-Gbx2+ ACs reach their mature dendritic morphology. To examine the dendritic morphology of individual S3-Gbx2+ ACs, we generated *Gbx2^CreERT2^*;*TIGRE^LSL-GFP^* (*Ai140D*) mice for single cell morphological analysis in retinal flat-mounts (Daigle et al., 2018; Kerstein et al., 2020). At P4 and P7, S3-Gbx2+ AC dendritic arbors were relatively small, symmetrical, and showed minimal change in arbor size (Fig. 1i-j, l-m). However, between P7 and P14, the dendritic arbors significantly increased in size and developed a marked asymmetry with dorsotemporal orientation (Fig 1k, l-n).

The dendrite growth and asymmetry between P7 and P14 coincides with eye opening in mice. During eye opening, some retinal neurons undergo light-dependent dendritic remodeling (Elias et al., 2018; El-Quessny et al., 2020; Tian and Copenhagen, 2003). Therefore, to test the hypothesis that light-dependent remodeling may regulate the dendritic morphology of S3-Gbx2+ ACs, we compared morphologies between light- and dark-reared animals. We observed no difference in dendritic morphology, symmetry, or orientation between the light- and dark-reared animals (Fig. S2). However, at P14, the dendrites from dark-reared animals had a small, but significant, increase in dendritic field area (Fig. S2d). These data show that eye opening and light-dependent developmental processes have no effect on S3-Gbx2+ AC dendrite morphology and only a small effect on arbor size.

### Embryonic deletion of *Gbx2* from AC precursors disrupts S3-Gbx2+ AC fate and dendrite lamination

Previous studies have demonstrated that *Gbx2* is an important transcription factor for neural development from neurogenesis to neuronal morphology in the brain and spinal cord (Chatterjee et al., 2012; Chen et al., 2010; Luu et al., 2011; Mallika et al., 2015). In the retina, the onset of *Gbx2* expression selectively in a subset of ACs from early postmitotic differentiation into adulthood suggests it may play a critical role in their specification and/or maturation (Kerstein et al., 2020). To determine the role of *Gbx2* in AC fate specification, we initially deleted *Gbx2* from AC precursors cells by generating *Ptf1a^Cre^*; *Gbx2^EGFP/Flox^* mice and *Ptf1a^Cre^*;*Gbx2^EGFP/+^* heterozygous littermates (Li et al., 2002); the *EGFP* transgene inserted into the endogenous *Gbx2* locus allowed us to genetically label and quantify the S3-Gbx2+ ACs. When *Gbx2* was deleted from AC precursors, we observed a significant reduction in the number of EGFP-positive cells in both the GCL and INL at P14 (Fig. 2a-f). Furthermore, in 2 out of 3 animals, we were unable to detect any EGFP-positive ACs in either layer (Fig. 2c, f). This phenotypic variability may be due to the transient expression of *Cre* driven from the *Ptf1a* locus in AC precursor cells, leading to incomplete deletion of *Gbx2* from the retina (Nakhai et al., 2007). Overall, these data suggest that in the absence of *Gbx2* expression, the S3-Gbx2+ ACs either fail to properly differentiate or have reduced survival in the developing retina.

**Figure 2.**
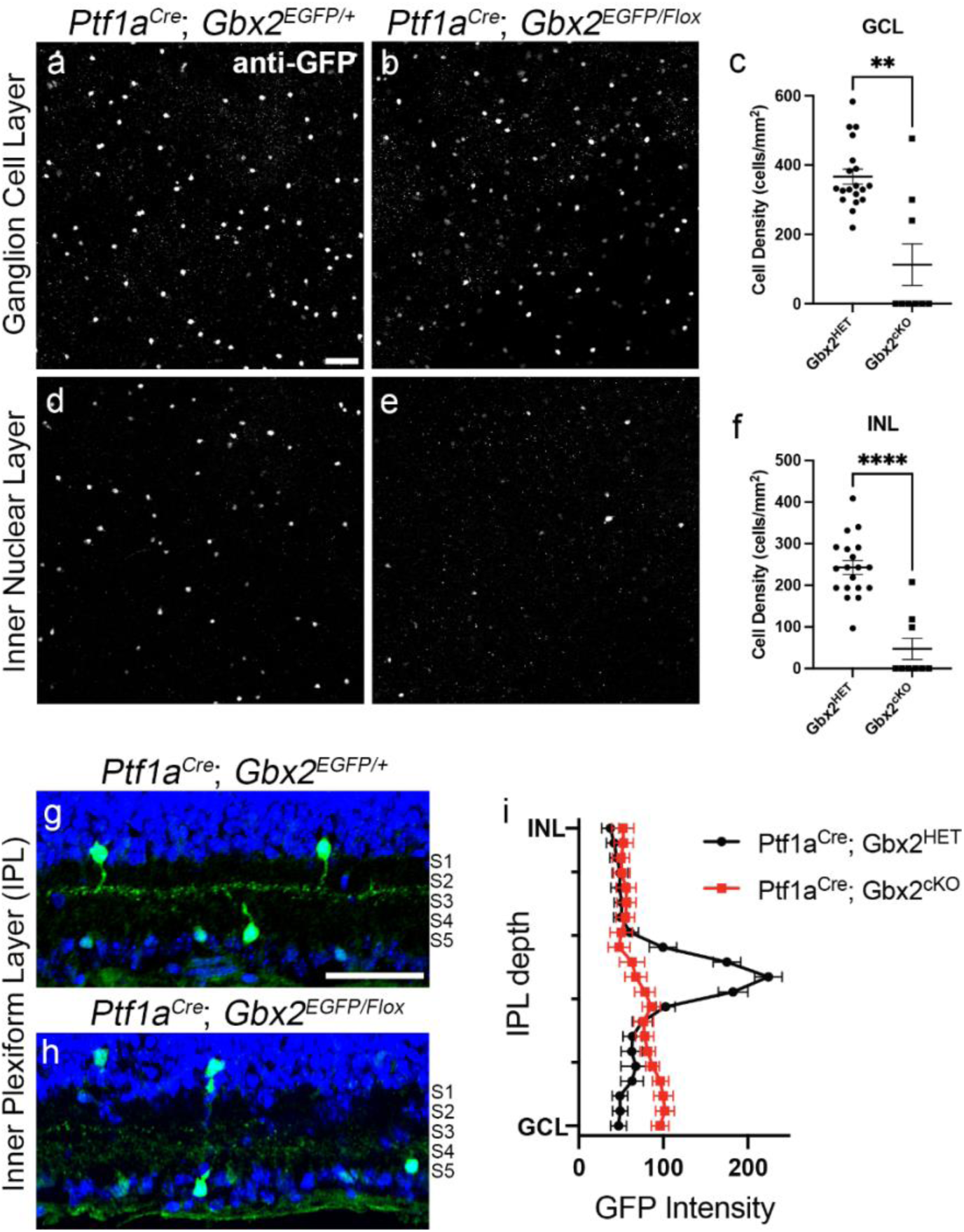
Embryonic deletion of *Gbx2* from AC precursors disrupts Gbx2+ AC cell fate and dendrite lamination. The number of Gbx2+ ACs was quantified in P14 retinal flat-mounts. EGFP-labels the cell bodies of Gbx2+ ACs in the ganglion cell layer (**a-b**) and the inner nuclear layer (**d-e**) in control *Ptf1a^Cre^; Gbx2^EGFP/+^* retinas (**a,d**) and mutant *Ptf1a^Cre^; Gbx2^EGFP/Flox^* retinas (**b, e**). Quantification of the cell body number in the ganglion cell layer (**c**) and inner nuclear layer (**f**). Adult retina sections from control *Ptf1a^Cre^; Gbx2^EGFP/+^* retinas (**g**) have EGFP+ dendrites that stratify in sublamina 3 (S3). In *Ptf1a^Cre^; Gbx2^EGFP/Flox^* mutants (**h**) Gbx2+ AC dendrites fail to stratify in S3. DAPI is in blue in **g** and **h**. A fluorescent intensity plot (**i**) shows the dendrite stratification for control (black) and *Ptf1a^Cre^; Gbx2^EGFP/Flox^* (red) retinas. **p=0.0026 and ****p<0.0001 using a Welch’s t-Test. Scale bar (**a, g**), 50μm.

We also examined if *Gbx2* contributes to proper morphological development of S3-Gbx2+ ACs. In control *Ptf1a^Cre^*; *Gbx2^EGFP/+^*retinas, S3-Gbx2+ ACs exhibit a precise, EGFP+ band of dendrites in S3 of the IPL at P14 (Fig. 2g, i). However, in *Ptf1a^Cre^*; *Gbx2^EGFP/Flox^* mutant retinas, the dendritic processes of the remaining EGFP+ cells are sparse and diffusely project throughout the IPL (Fig. 2h-i). These data show that *Gbx2* not only controls cell specification, but also that it is likely required for the proper targeting of S3-Gbx2+ AC dendrites to the correct layer in the IPL.

### Early, but not late, postnatal deletion of *Gbx2* disrupts dendrite stratification

To better understand how Gbx2 regulates dendrite stratification in the retina, we developed a genetic strategy to delete *Gbx2* after cells are specified. *Gbx2^CreERT2/Flox^*; *Rosa26^LSL-TdTomato^* mutants and *Gbx2^CreERT2/+^*;*Rosa26^LSL-TdTomato^* heterozygous littermate controls were dosed with tamoxifen (60 mg/kg) at P1 to delete *Gbx2* at the earliest stages of dendrite development (Fig 1a), while simultaneously genetically labeling cells with TdTomato. This approach labels two AC types: the S3-stratifying subtype that expresses high levels of *Gbx2*, and a second subtype that expresses low levels of Gbx2 and stratifies in S5 (Fig. 3a) (Kerstein et al., 2020). When *Gbx2* was deleted at P1, we consistently observed ectopic TdTomato+ dendritic projections stratifying in the upper layers of the IPL at the border of S1 and the INL (Fig. 3a-b, e). In contrast, when we deleted *Gbx2* after retina development was complete (60 mg/kg tamoxifen at P35), we did not observe any ectopic projections (Fig. 3c-d). Therefore, Gbx2 is required for the proper dendrite stratification in the IPL during development but is not required to maintain this stratification in adulthood.

**Figure 3.**
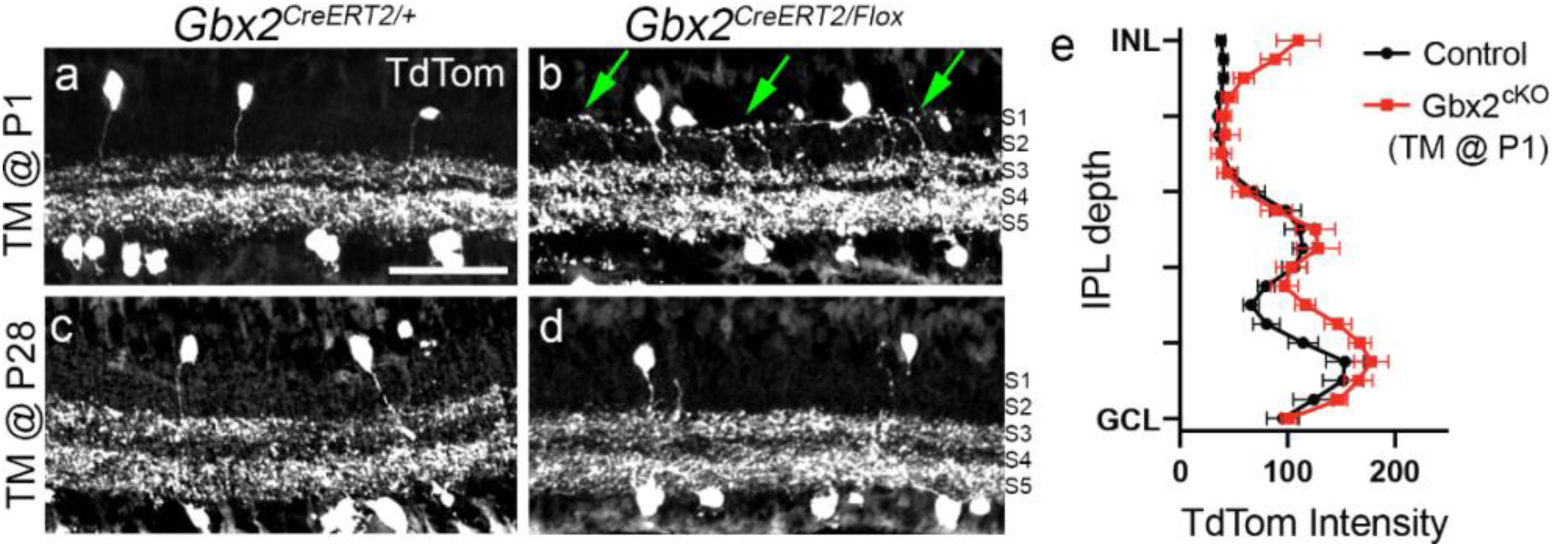
Early (P1), but not late (P28), postnatal deletion of *Gbx2* disrupts dendrite lamination. Adult retinal sections were labeled for Gbx2+ ACs (TdTomato). Adult retina sections from (**a**) control Gbx2+ ACs (Gbx2^CreERT2/+^) and (**b**) conditional knockout of *Gbx2* from the Gbx2+ ACs (*Gbx2^CreERT2/Flox^*) when mice were administered tamoxifen (0.025 mg, oral gavage) at P1. Arrows denote ectopic dendrite projections into S1. Adult retina sections from (**c**) control Gbx2+ ACs (Gbx2^CreERT2/+^) and (**d**) conditional knockout of *Gbx2* from the Gbx2+ ACs (*Gbx2^CreERT2/Flox^*) when mice were administered tamoxifen (1 mg, oral gavage) at P28. (**e**) Quantification of dendrite fluorescence intensity (TdTomato) by IPL depth from control (black) and *Gbx2^CreERT2/Flox^* (red) retinas. Scale bar, 50μm.

### *Gbx2* effector genes *Robo1* and *Robo2* are expressed in the developing retina

*Gbx2* functions cell autonomously to regulate the proper dendrite stratification of Gbx2+ ACs. In developing thalamic neurons, Gbx2 directly regulates the expression of *Robo1* and *Robo2*, which have well-established roles in axon guidance and neuronal morphology (Chatterjee et al., 2012; López-Bendito et al., 2007; Mallika et al., 2015). We examined *Robo1* and *Robo2* expression in the retina at P7, when Gbx2 dendrites are actively stratifying in S3. Consistent with scRNAseq datasets of different retinal neuron populations (Yan et al., 2020), *Robo1* and *Robo2* were expressed in ACs and retinal ganglion cells (RGCs) within INL and GCL (Fig. 4a-b, arrowheads). Using immunohistochemistry, we found that Robo1 and Robo2 receptors were enriched in AC and RGC dendrites in the IPL and RGC axons in the GCL (Fig. 4c-d). In addition, our previous RNAseq dataset from purified Gbx2+ ACs showed expression of both *Robo1* and *Robo2* (Kerstein et al., 2020).

**Figure 4.**
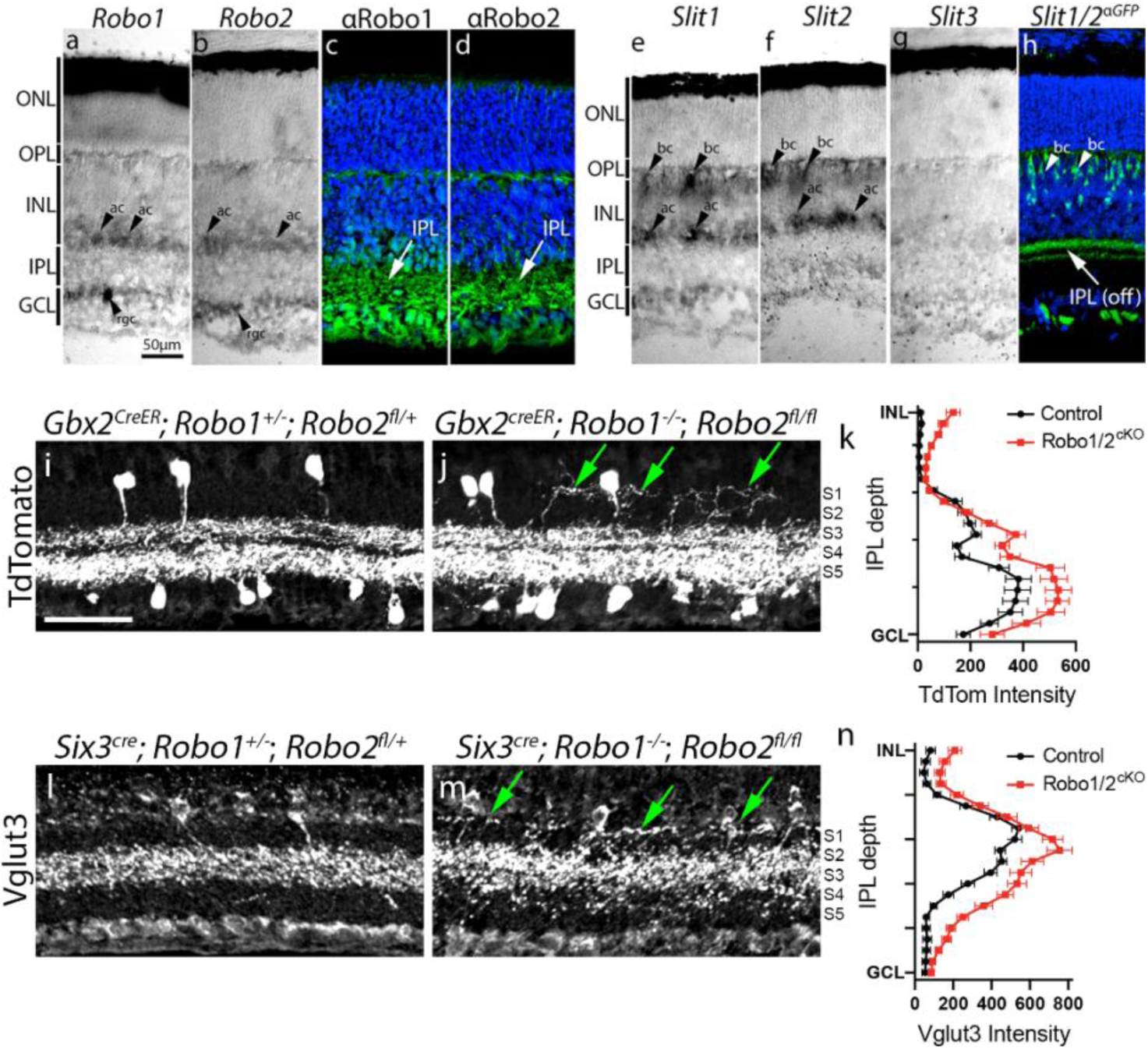
Deletion of *Robo1/2* disrupts dendrite lamination of multiple AC subtypes. *In situ* hybridization for (**a**) *Robo1* and (**b**) *Robo2* expression in P8 retinal sections. Immunolabeling for (**c**) Robo1 and (**d**) Robo2 localization; arrows denote Robo localization to neuronal processes in the IPL. *In situ* hybridization for (**e**) *Slit1*, (**f**) *Slit2*, and (**g**) *Slit3* expression in P8 retinal sections. (**h**) *Slit1/2* expressing cells are marked with transgenic GFP expression labeling in P7 retinal sections from *Slit1^Tau-GFP/+^; Slit2^Tau-GFP^* mice. Arrow denotes *Slit* expressing processes in OFF layers of IPL. Arrowheads: ac, amacrine cell; bc, bipolar cell; rgc, retinal ganglion cell. (**i-j**) Adult retinal sections were labeled for Gbx2+ ACs (TdTomato) from (**i**) control and (**j**) *Gbx2^CreERT2^; Robo1^-/-^; Robo2^Flox/Flox^* mutant retinas. Tamoxifen (0.025 mg, oral gavage) was administered at P1. (**k**) The fluorescent intensity plot shows the dendrite stratification (TdTom) for control (black) and *Gbx2^CreERT2^; Robo1^-/-^; Robo2^Flox/Flox^* (red) retinas. (**l-m**) Adult retinal sections immunolabeled for Vglut3 from (**l**) control and (**m**) *Six3^Cre^; Robo1^-/-^; Robo2^Flox/Flox^*mice. (**n**) The fluorescent intensity plot shows the dendrite stratification (TdTom) for control (black) and *Six3^Cre^; Robo1^-/-^; Robo2^Flox/Flox^* (red) retinas. Green arrows point to ectopic dendrite projections into the S1 of the IPL. Scale bar, 50μm.

Robo1 and Robo2 bind secreted Slit ligands to induce axon repulsion in several regions of the developing nervous system, including the retina (Brose et al., 1999; Kidd et al., 1999). *Slit1* and *Slit2* were both expressed in subsets of bipolar cells and ACs of the developing retina (Fig. 4e-f, arrowheads). *Slit3* exhibited little to no expression, consistent with previous studies (Fig. 4g) (Erskine et al., 2000; Rama et al., 2015). In scRNAseq datasets, *Slit1* and *Slit2* are selectively expressed in OFF layer bipolar cells (Types 1a, 1b, 2, 3a) (Shekhar et al., 2016). Using a *Slit1^EGFP/+^;Slit2^EGFP/+^*mouse (Plump et al., 2002), we found that *Slit1/2*-expressing bipolar cells stratify their axons selectively in the OFF layers of the IPL (S1-2) (Fig. 4h). These data show that bipolar cell-derived Slit ligands are in the right location at the right time to form a repulsive barrier in the OFF layers of the IPL and may contribute to the segregation of ON-OFF retinal circuitry.

### Deletion of *Robo1* and *Robo2* receptors disrupts AC dendrite lamination

Once we determined that both Robo receptors and Slit ligands are both present in the developing IPL, we tested whether they regulate AC dendrite lamination. We generated *Gbx2^CreERT2^*; *Robo1^-/-^*; *Robo2^Flox/Flox^*; *Rosa26^LSL-TdTomato^* mice, and administered Tamoxifen at P1 (60 mg/kg) to delete Robo receptors from Gbx2+ ACs after their differentiation, but before dendrite development. In control *Gbx2^CreERT2^*; *Robo1^-/+^*; *Robo2^Flox/+^*; *Rosa26^LSL-TdTomato^* mice, TdTomato+ dendrites stratified normally in S3 and S5. In contrast, when we deleted *Robo1/2* receptors from Gbx2+ ACs, the development of their dendrites was disrupted, with a significant increase in ectopic projections into the OFF layers of the IPL (Fig. 4i-j). When we analyzed the fluorescence intensity of TdTomato+ dendrites in *Gbx2^CreERT2^*; *Robo1^-/-^*; *Robo2^Flox/Flox^*; *Rosa26^LSL-TdTomato^* mice across the IPL, we noticed two changes in the *Robo1/2* mutant retinas: a significant increase of dendrite projections into S1, and an overall increase in fluorescence intensity in the S3 and S5 layers (Fig. 4K). These data show that Robo receptors function cell autonomously to control dendrite lamination in the Gbx2+ ACs.

Next, we determined if Robo receptors were important for controlling the stratification of other AC subtypes. To test this, we deleted *Robo1* and *Robo2* receptors from retinal progenitor cells using a *Six3^Cre^*; *Robo1^-/-^*; *Robo2^Flox/Flox^*mouse. Based on the specific antibody immunolabeling, the deletion of *Robo1/2* receptors from the developing retina did not alter the stratification or morphology of several bipolar cell subtypes, suggesting *Slit1/2* expressing cells still correctly localized their axon terminals to the OFF layers of the IPL (Fig. S3). However, we found that another S3-targeting AC, the Vglut3+ AC, also mistargets its dendrites to the OFF layers (S1) of the IPL in *Six3^Cre^*; *Robo1^-/-^*; *Robo2^Flox/Flox^* retinas (Fig. 4l-n). Like the Gbx2+ ACs, Vglut3+ ACs showed an overall increase in dendrite density in Robo1/2 mutant retinas (Fig. 4m-n). In contrast, when we deleted Robo1/2 receptors selectively from Gbx2+ ACs (*Gbx2^CreERT2^*; *Robo1^-/-^*; *Robo2^Flox/Flox^*; *Rosa26^LSL-TdTomato^*), we did not observe dendritic defects in the Vglut3+ ACs (Fig. S4a-c). In contrast to the Gbx2+ ACs and Vglut3+ ACs that stratify in S3, we observed no change in starburst AC dendrite stratifications in S2 and S4 in the *Robo1/2* mutant retinas, suggesting that the abnormal stratification in *Six3^Cre^*; *Robo1^-/-^*; *Robo2^Flox/Flox^* mice was selective for specific AC subtypes (Fig. S4e-f). Thus, in addition to Robo receptors having an important role in Gbx2+ AC development, it appears they also have a broader role in dendrite lamination in other AC subtypes, preventing some ON layer dendrites from targeting OFF layers of the IPL.

## DISCUSSION

During development, the mouse retina generates >120 molecularly distinct neuronal subtypes. About half of this cellular diversity comes from the amacrine cells (ACs). The generation of AC subtypes—each with their own unique morphology and function—requires specific genetic signaling pathways. Here, we focused on how the transcription factor *Gbx2* and its effector genes control the cell fate and development of a single AC subtype. We found the majority of S3-Gbx2+ ACs are born and acquire their cell fate between E14-E18 (Fig. S1). These cells stratify their dendrites in S3 of the inner plexiform layer (IPL) over the first week of postnatal development, and subsequently develop their characteristic asymmetric morphology during the second postnatal week (Fig. 1). Deletion of *Gbx2* from AC precursor cells significantly reduces the number of ACs that acquire the Gbx2+ AC fate (Fig. 2). This is similar to previous studies that have shown that the transcription factors *Isl1* and *Lhx9* are important for Starburst (ChAT+) and Nitric Oxide Synthase (NOS+) AC fates, respectively (Balasubramanian et al., 2018; Elshatory et al., 2007). *Gbx2* continues to play an important role at subsequent stages of development by controlling dendrite lamination of Gbx2+ ACs. Deletion of *Gbx2* from the Gbx2+ ACs after differentiation but before dendrite formation leads to ectopic projections within the IPL (Fig. 3). This role of *Gbx2* in dendrite targeting is analogous to the role that other transcription factors—such as *Sox2*, *Lhx9*, and *Fezf1*—play in AC migration and dendrite targeting (Balasubramanian et al., 2018; Peng et al., 2020; Stincic et al., 2018; Whitney et al., 2014). Dendrite targeting in the Gbx2+ AC is controlled at least in part by the transmembrane receptors Robo1 and Robo2. Deletion of *Robo1* and *Robo2* from Gbx2+ ACs phenocopied the ectopic dendrites in the *Gbx2* mutant retinas (Fig. 4). We also showed that Robo receptors are necessary for dendrite targeting in other S3-targeting AC subtypes, such as the Vglut3+ ACs (Fig. 4). Based on the location of Slit ligands and the ectopic projections observed in *Robo1/2* mutant retinas, we hypothesize that *Slit1/2* may act as a repulsive barrier in the OFF layers of the IPL to prevent *Robo1/2*-expressing ON layer dendrites from extending into the wrong layers. Therefore, Slit-Robo signaling is a mechanism that contributes to the organization of ON and OFF pathways in the developing retina.

In the developing brain, *Gbx2* is a critical transcription factor for neuronal specification, migration, and connectivity. Depending on the timing and tissue, the deletion of *Gbx2* leads to the disruption of several developmental processes: the generation and specification of thalamic neurons, the patterning of spinal interneurons, the migration of cholinergic striatal neurons, and the proper guidance of thalamocortical axons (Chatterjee et al., 2012; Chen et al., 2010; Luu et al., 2011; Mallika et al., 2015). In this study, we investigated the role of Gbx2 on retinal AC specification and dendrite targeting. Using multiple *Gbx2* cKO mice, we show that *Gbx2* is required for both the initial subtype specification and subsequent dendrite targeting of Gbx2+ ACs. We also observed the same dendritic mis-projections when we deleted the *Gbx2* effector genes, *Robo1* and *Robo2*, from the Gbx2+ ACs (Fig. 4). The deletion of *Gbx2* from thalamic neurons or migrating neural crest cells alters the expression of Robo receptors (Mallika et al., 2015; Roeseler et al., 2012). However, it was not clear whether this is due to direct or indirect regulation of *Robo1* and *Robo2* expression by *Gbx2*, or due to altered fate of the *Gbx2^cKO^* thalamic neurons. *In vitro* studies using ChIPseq in human cell lines have shown Gbx2 can directly bind to the *ROBO1* gene, suggesting it may directly regulate its expression (Roeseler et al., 2012). However, it is unknown if *Gbx2* directly regulates the expression of *Robo* receptors in primary neurons. Unfortunately, due to the lack of a ChIP-grade antibody for *Gbx2* and the relatively small number of Gbx2+ ACs in the retina, we were unable to test the binding of *Gbx2* to *Robo1* and *Robo2* genes in retinal neurons. Therefore, we cannot rule out that *Gbx2* may regulate Robo receptor expression indirectly through other transcription factors. For example, *Gbx2* is thought to regulate Robo receptor expression indirectly through the transcription factors *Lmo3* and *Lhx9* in thalamic neurons (Chatterjee et al., 2012). *Lhx9* is expressed in multiple AC subtypes, including the Gbx2+ ACs (Kerstein et al., 2020; Yan et al., 2020). Like *Gbx2*, *Lhx9* is expressed in ACs that stratify in the S3 and S5 sublamina in the IPL (Balasubramanian et al., 2014a, 2014b). When *Lhx9* is deleted from the retina, dendrites of Lhx9+ ACs mis-project into the upper layers of IPL, similar to what we observed in the *Gbx2* and Robo1/2 cKOs (Balasubramanian et al., 2018). Therefore, *Gbx2* may regulate the expression of *Robo* receptors through direct binding or indirectly through expression of *Lhx9*.

Robo1 and Robo2 receptors bind to the extracellular Slit ligands to induce repulsion during axon outgrowth and cell migration (reviewed in (Blockus and Chédotal, 2016, 2014)). However, increasing evidence suggest Slit-Robo signaling plays additional roles during neuronal development, such as neuronal proliferation, dendrite branching, and synapse formation (Blockus et al., 2021; Borrell et al., 2012; Gibson et al., 2014; Whitford et al., 2002). Slit1/2 and Robo1/2 are expressed within the retina, but the role of Slit-Robo interactions in dendrite stratification within the retina is poorly defined (Fig 4) (Erskine et al., 2000; Rama et al., 2015). Slit-Robo signaling restricts RGC axons to the nerve fiber layer as they extend towards and exit the optic nerve head (O’Sullivan et al., 2017; Thompson et al., 2006). In addition, the incoming vasculature requires Slit-Robo signaling as it expands across the retina (Rama et al., 2015). Here, we show that Slit1/2-expressing bipolar cells stratify their axons in the OFF layers of IPL (S1-S2) (Fig. 4h). These OFF layers are the precise location where ectopic projections from Robo1/2 expressing S3 stratifying ACs (Gbx2+ and Vglut3+) misproject in *Gbx2* and *Robo1/2* cKO retinas. Thus, while indirect, this result provides support to the model that bipolar cell derived Slits create a repulsive boundary in the OFF layers to prevent the ectopic stratification of Robo+ dendrites. Not all AC subtypes require Slit-Robo signaling, as OFF SACs stratify normally in S2 in *Robo1/2* cKO retinas. Other ligand-receptor pairs are important for the reciprocal interaction, with the transmembrane ligand *Sema6a* preventing stratification of dendrites expressing the semaphorin receptor *PlexinA4* in ON layers of IPL (Matsuoka et al., 2011). These previous findings in combination with our results suggest that both Sema6a-PlexinA4 and Slit1/2-Robo1/2 signaling are necessary for ON-OFF circuit formation during retina development.

One limitation of this current study is that we do not know how consequences of deleting *Gbx2* or *Robo1/2* from the retina disrupts visual function. Our previous study demonstrated that Gbx2+ ACs receive excitatory inputs from both ON and OFF bipolar cells (Kerstein et al., 2020). One question our results raise is whether the ectopic dendritic projections of the Gbx2+ ACs from *Gbx2* or *Robo1/2* mutant retinas would potentially increase inputs from OFF bipolar cells. However, as we do not currently know the identity of the downstream targets of Gbx2+ ACs, it is difficult to assess how either the changes in AC specification or the mis-targeting of dendrites observed in the *Gbx2* mutants would affect visual behaviors in the mouse. However, the abnormal dendrite targeting by Vglut3+ ACs in *Robo1/2* cKOs can be assessed in future studies. Vglut3+ ACs serve an important role in the detection of object motion and visual threats, and defects in Vglut3+ AC synapse formation cause deficits in differential motion sensing (Kim et al., 2020; Kim and Kerschensteiner, 2017; Krishnaswamy et al., 2015). Therefore, the mistargeting of Vglut3+ AC dendrites we observed in the *Robo1/2* mutant retinas may also reduce connectivity and result in a loss of object motion detection.

Our findings are potentially important for understanding visual disorders such as dyslexia. Polymorphisms in the human *ROBO1* gene and promotor regions are associated with dyslexia (Hannula-Jouppi et al., 2005; Massinen et al., 2016). While the etiology of dyslexia is thought to be due to changes in cortical connectivity, it is possible that changes in retinal circuitry like those described in this study may also contribute to the visual disruptions of the disease. Ultimately, identifying the genetic mechanisms of neuronal subtype specification and neural circuit assembly will increase our understanding of how genes lead to behaviors and what molecular pathways are involved in human disease.

## METHODS AND MATERIALS

### Key Resources Table

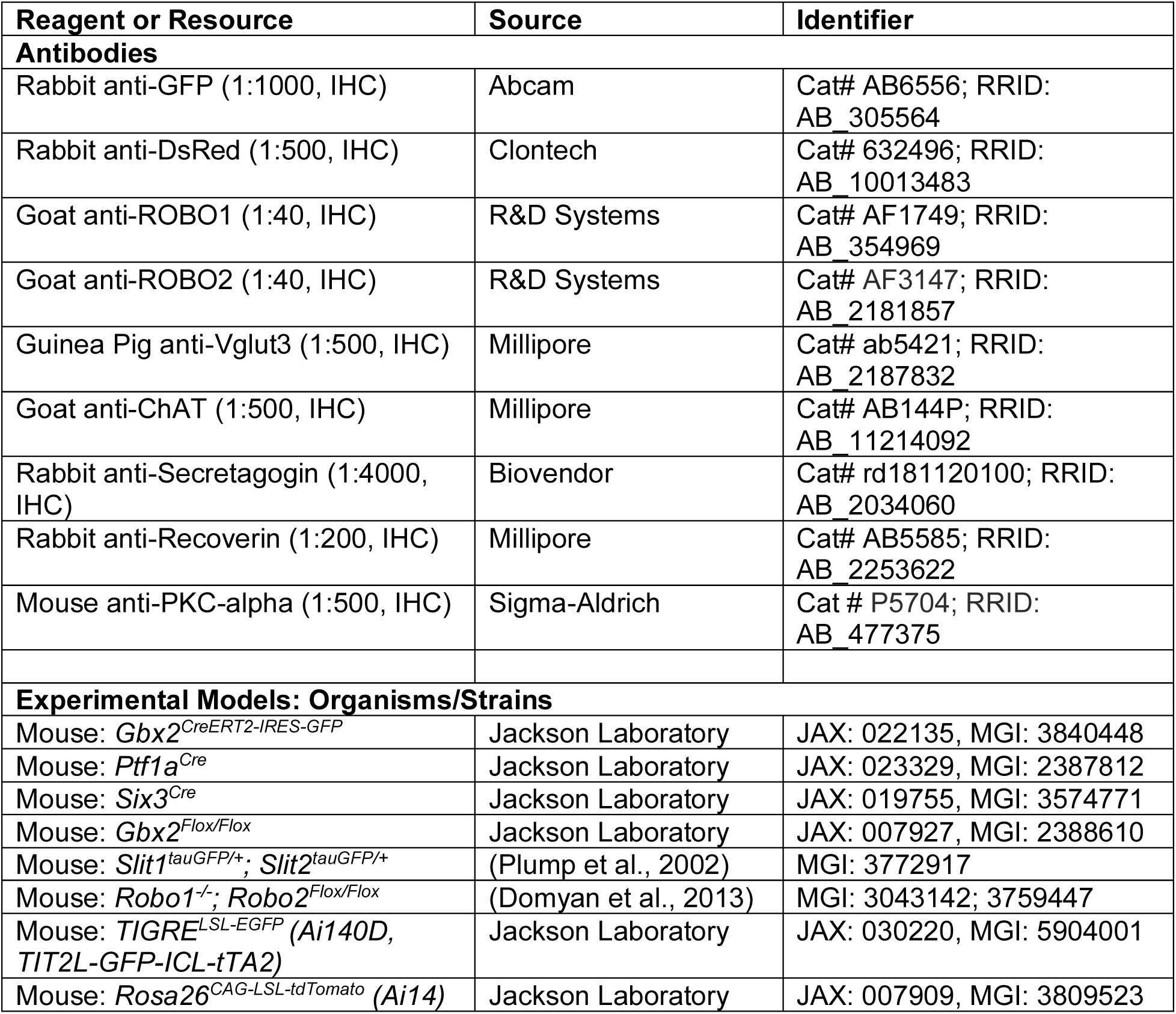

### Resource Availability

#### Lead Contact

Requests for additional information and resources should be directed to the Lead Contact, Patrick Kerstein.

#### Materials Availability

No mice or reagents were generated in this study. The mouse lines used in the study will be provided with permission from the investigator that generated them (see Key Resource Table). Question and inquiries should be direct to the Lead Contact.

#### Data and Code Availability

All data generated in this manuscript is available on request. Please contact the Lead Contact. No new code was generated in this study.

### Experimental Model and Subject Details

#### Animals and Animal Procedures

All experiments in this study used mice from both sexes on a mixed genetic background. The following Cre mouse lines were used for tissue-specific fluorescent labeling and gene deletion: *Gbx2^CreERT2-IRES-EGFP^*(Chen et al., 2009), *Ptf1a^Cre^* (Nakhai et al., 2007), and *Six3^Cre^*(Furuta et al., 2000). The following knock-in, knockout, and conditional knockout mouse lines were used: *Gbx2^Flox/Flox^* (Li et al., 2002), *Robo1^-/-^*; *Robo2^Flox/Flox^* (Domyan et al., 2013; Long et al., 2004; Lu et al., 2007; Ma and Tessier-Lavigne, 2007), and *Slit1^tauGFP/+^; Slit2^tauGFP/+^* (Plump et al., 2002). Two different Cre-dependent reporter mouse lines were used: Ai140D/*TIGRE^TRE2-LSL-GFP; CAG-LSL-tTA2^* (Daigle et al., 2018) and Ai14/*Rosa26^LSL-TdTomato^* (Madisen et al., 2010). In experiments that used the *Gbx2^CreERT2^* mouse line, tamoxifen dissolved in sunflower seed oil was administered in P1 or P28 mice. At the P1 timepoint, 50µl of 0.5mg/mL tamoxifen was injected into the milk pouch of mouse pups as previously described (Pitulescu et al., 2010). For sparse labeling, 50 µl of 0.01mg/mL tamoxifen was injected. At the P28 timepoint, 200µl of 5mg/mL was administered by oral gavage. For dark-rearing experiments, neonatal mice (P0-P14) were housed with their nursing mothers in an environmental chamber (Percival Scientific, Perry, IA) maintained at 20-21°C on a 24 hour dark cycle, with free access to food and water. All animal procedures were approved by Oregon Health and Science University Institutional Animal Care and Use Committee, the Purdue Animal and Use Committee (PACUC) at Purdue University, and IACUC of Thomas Jefferson University and conformed to the National Institutes of Health’s *Guide for the Care and Use of Laboratory Animals*.

### Method Details

#### Immunohistochemistry and *in situ* hybridization

Neonatal and adult retinas were prepared for immunolabeling by removing the eyes from euthanized animals. For tissue fixation, the cornea was removed, and the eyes were placed in 4% EM-grade paraformaldehyde (PFA) for 30 mins at room temperature. Next, the eyes were washed twice in 1X PBS for 30 mins. For cryo-preservation, eyes were placed in 15% sucrose for at least 2 hours at 4°C. The lens was then removed, eyes were embedded in a cryo-mold in Optimal Cutting Temperature media and frozen. 20µm retinal sections were collected on a glass microscope slide (Fisher Superfrost Plus) using a cryostat. For immunolabeling, slide-mounted retinal sections were washed with 1X PBS for 10 mins, and then blocked with 2% normal donkey serum (NDS) with 0.2% Triton X-100 for 30 mins. Retinal sections were incubated with primary antibodies in NDS blocking buffer overnight at 4°C. The primary antibodies and their dilutions used in this study are listed in the Key Resources Table. Next, sections were washed in 3 times in NDS blocking buffer for 10 mins, and incubated in secondary antibodies in blocking buffer for 2 hours at room temperature. Retinal sections were washed three times in PBS for 10 mins with either Hoechst or DAPI (1:5000) stain included in the first wash step. Tissue was mounted in Fluoromount-G (Southern Biotech) and cover-slipped for imaging.

Retinal flat-mounts were dissected and washed as described above. The retinas were removed from the eye cup and flattened by making 3-4 equally spaced incisions into the center of the tissue. Retina flat-mounts were post-fixed in 4% PFA for 30 mins to help maintain their flat shape. Retinas were washed 3 times in 2% NDS and 0.2% Triton X-100 for 30 mins at room temperature. Retinas were incubated with primary antibodies in blocking solution for 3-7 days at 4°C. After incubation with primary antibody, the retinas were washed 3 times with blocking solution for 30 mins. Next, retinas were incubated with secondary antibody in blocking solution overnight at 4°C. Retinas were then washed three times in PBS for 30 mins with either Hoechst or DAPI (1:5000) stain included in the first wash step. Tissue was mounted on a slide with Fluoromount-G and cover-sliped for imaging.

For *in situ* hybridization analysis, retinas were dissected from P8 mouse pups, fixed in 4% paraformaldehype, embedded in optimal cutting temperature medium and cryosectioned at 20 µm. Digoxigenin-labeled riboprobes were generated from pBluescript vectors for *Robo1*, *Robo2, Slit1*, *Slit2*, and *Slit3*, as previously developed and used in the mouse retina (Erskine et al., 2000; Yuan et al., 1999).

#### Fluorescence Image Acquisition

All retinal sections were imaged on a Zeiss Axio Imager M2 upright microscope equipped with an ApoTome2 system using a 20x objective. Retinal flatmounts for single cell morphology, dendrite orientation, and cell density assays were imaged on a Zeiss LSM 900 confocal microscope using a 20x objection. All images were acquired using the Zeiss Zen Imaging software on both microscope setups.

### Quantification and Statistical Analysis

#### Neuronal Morphology and Density Analysis

All analysis of neuronal morphology and cell density was completed off-line using Imaris (Bitplane) or FIJI software (Schindelin et al., 2012). For dendrite stratification in retinal cross-sections, measurements of fluorescent intensity were made along the IPL depth using the FIJI plugin IPLaminator (Li et al., 2016). All values were binned into 5% increments along the IPL depth. For dendritic arbor morphology analysis in retinal flatmounts, we analyzed isolated single cells sparse labeled in the *Gbx2^CreERT2-IRES-EGFP^*; *Tigre^LSL-EGFP^* mice. Tracing and analysis of dendritic arbors were completed using the Filaments plugin in Imaris. Cell density of EGFP+ cells was calculated by the mean of 4 measurements from each animal in the INL and GCL. All cell density measurements were made at least 200 µm from the optic nerve head and peripheral retinal edge. Symmetry index was calculated by subtracting the angles of missing dendrite from 360 and then divided by 360, as previously described (Kerstein et al., 2020; Sun et al., 2013). Dendrite orientation was calculated by the direction of the longest dendrite branch and then measurements were binned into 8 different groups based on the cardinal directions.

#### Statistics

In these analyses, each experiment and timepoint included measurements from a minimum of 3 retinas from at least 3 different animals. For all datasets, the variance was reported as the mean ± SEM. Each dataset was tested for a normal distribution with a D’Agostino & Pearson test. Analysis between two groups was completed using an unpaired student’s t-test (parametric) or a Mann-Whitney U test (nonparametric). For analysis between more than two groups, we used a one-way analysis of variance (ANOVA) with a Tukey’s multiple comparison test (parametric) or a Kruskal-Wallis with a Dunn’s multiple comparison test (nonparametric). For IPL lamination assays, groups and individual IPL depths were statistically test using a 2-way ANOVA with multiple comparisons test. All statistical tests were performed using Prism 9 software (Graphpad Software, Inc.).

## ACKNOWLEDGEMENTS

We would like to thank the members of the Wright laboratory for their assistance and discussion throughout the course of this study. We also thank Dr. Charles Allen (OHSU) for use of his temperature- and light-controlled environmental chambers for dark rearing. This work was supported by NIH grant R01EY032057 and R01EY032057-S to K.M.W.; NIH grant R01NS112504 to L.M.; NIH grant F32EY029974, the Collins Medical Trust, and the Knights Templar Eye Foundation to P.C.K. Confocal microscopy and analysis was performed in the OHSU Advance Light Microscopy Core supported by the NIH grant P30 NS061800.

## FIGURE LEGENDS

**Supplemental Figure 1 (S1).**
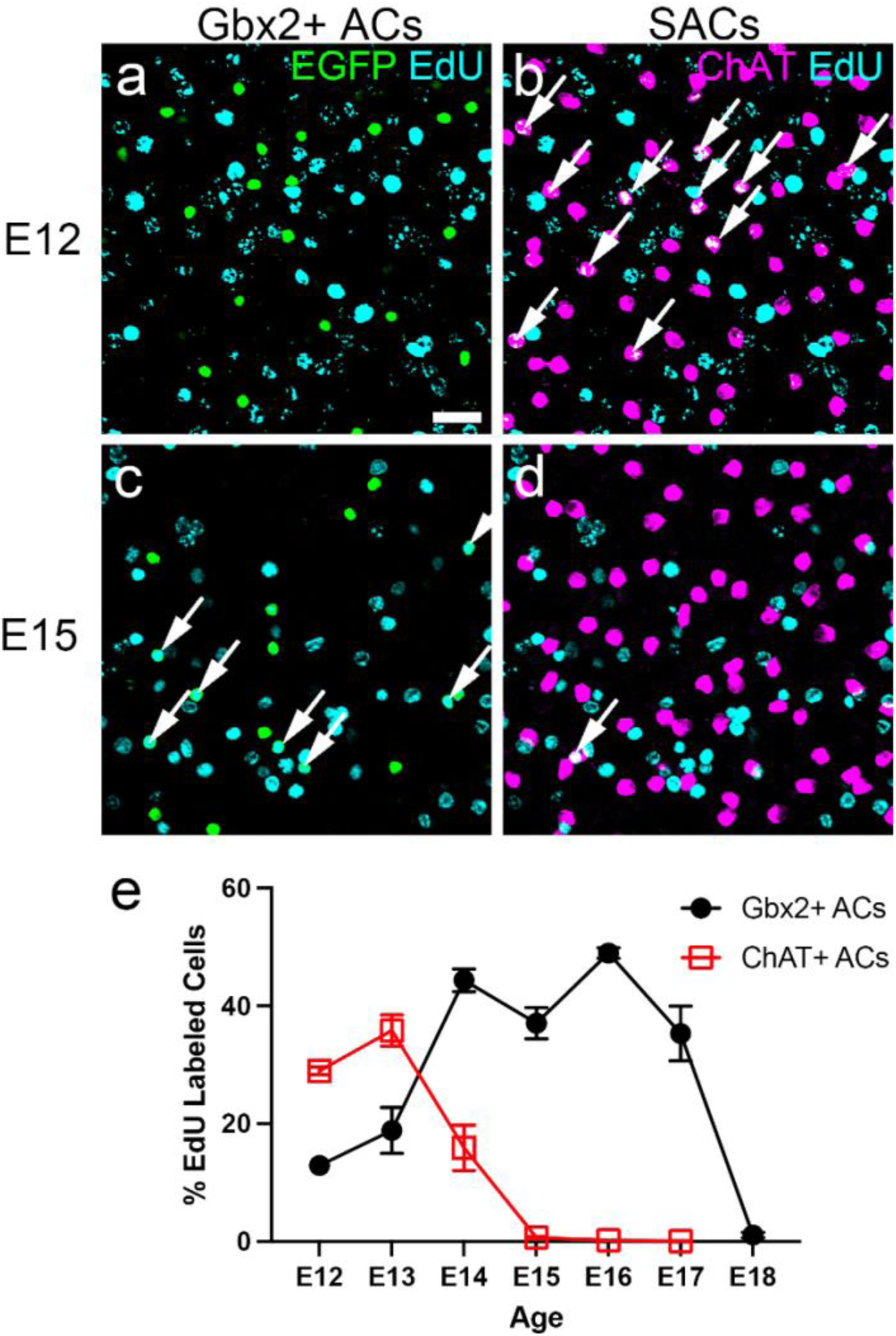
Birthdating of Gbx2+ amacrine cell proliferation using EdU labeling. (**a-b**) Co-localization of EdU-birthdated cells (cyan) at E12 with (**a**) the Gbx2+ AC marker EGFP (green) and (**b**) the SAC marker ChAT (magenta) in the GCL of adult retinas from the *Gbx2^EGFP^* mouse. (**c-d**) EdU-birthdated cells (cyan) at E15 co-labeled with markers for (**c**) Gbx2+ ACs (green) and (**d**) SACs (magenta) in the GCL of the adult retina. Arrows denote co-localization of EdU with the AC subtype marker. (**e**) Distributions are quantified as percent of EdU+ cells over the total labeled cells by each marker. For each timepoint, n=3-8 animals. Scale bar, 25 µm.

**Supplemental Figure 2 (S2).**
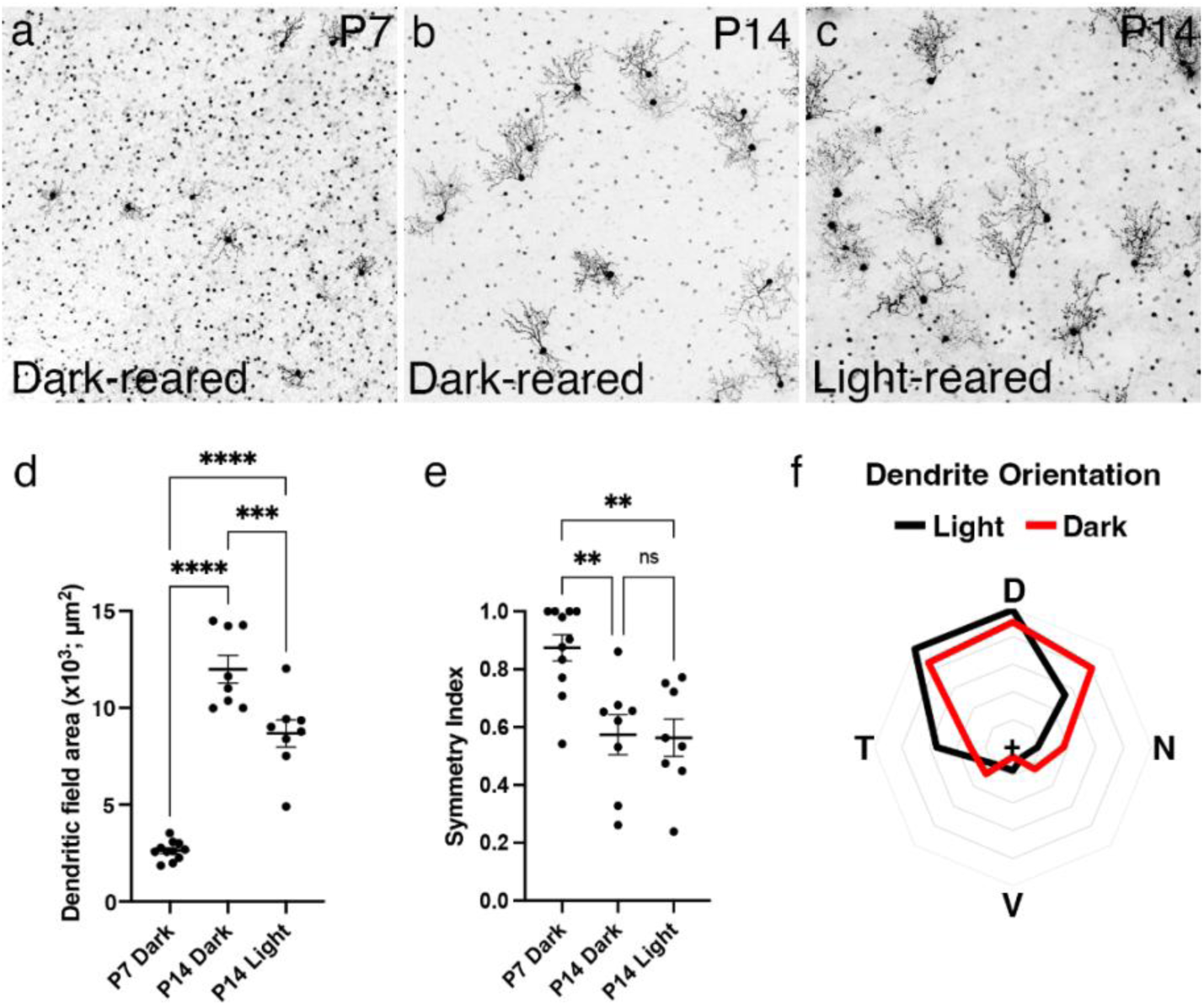
Dendrite morphology in S3-Gbx2+ ACs does not develop using a light-dependent mechanism. Dendrite morphology of S3-Gbx2+ ACs *en face* from (**a**) P7 and (**b**) P14 of dark-reared mice; and from (**c**) P14 control (light-reared) mice. Gbx2+ ACs are sparely labeled using the *Gbx2^CreERT2^*; *TIGRE^LSL-EGFP^* mouse line and low dose of tamoxifen (0.01mg) at P1. Quantification of Gbx2+ AC dendritic arbor (**d**) area and (**e**) symmetry at P7 and P14 developmental timepoints. (**f**) Polar plot showing the asymmetric orientation of S3-Gbx2+ AC dendrites in dorsal-temporal direction at P14 in light reared animals (black) is preserved in dark-reared animals (red). In (**d**) and (**e**), ns, not significant, **p<0.01, ***p<0.001, and ****p<0.0001 using a One-way ANOVA with multiple comparisons test. Scale bar, 50μm.

**Supplemental Figure 3 (S3).**
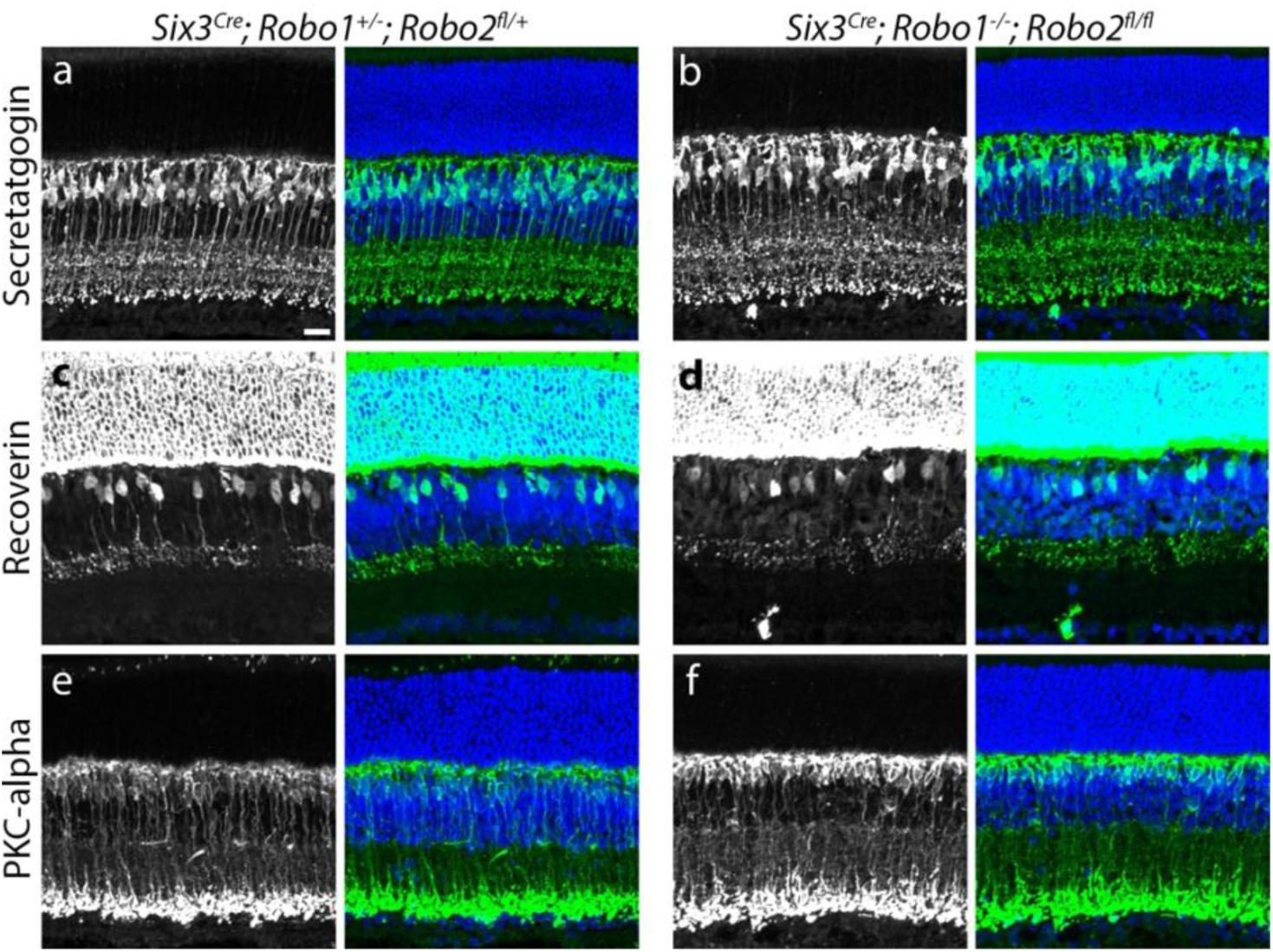
The pan-retinal deletion of *Robo1* and *Robo2* does not disrupt bipolar cell axon stratification. In adult retinal sections, the axon stratification of bipolar cell subtypes was compared between the (**b, d, f**) *Six3^Cre^; Robo1^-/-^; Robo2^Flox/Flox^* conditional knockouts and (**a, c, e**) their heterozygous littermate controls. Using immunohistochemistry, the following bipolar cell subtypes were compared: (**a, b**) types 2, 3, 4, 5, 6, and 8 cone bipolar cells by secretagogin, (**c, d**) types 1 and 2 OFF cone bipolar cells by Recoverin, and (**e, f**) rod bipolar cells by PKC⍺. Scale bar, 25μm in (**a**).

**Supplemental Figure 4 (S4).**
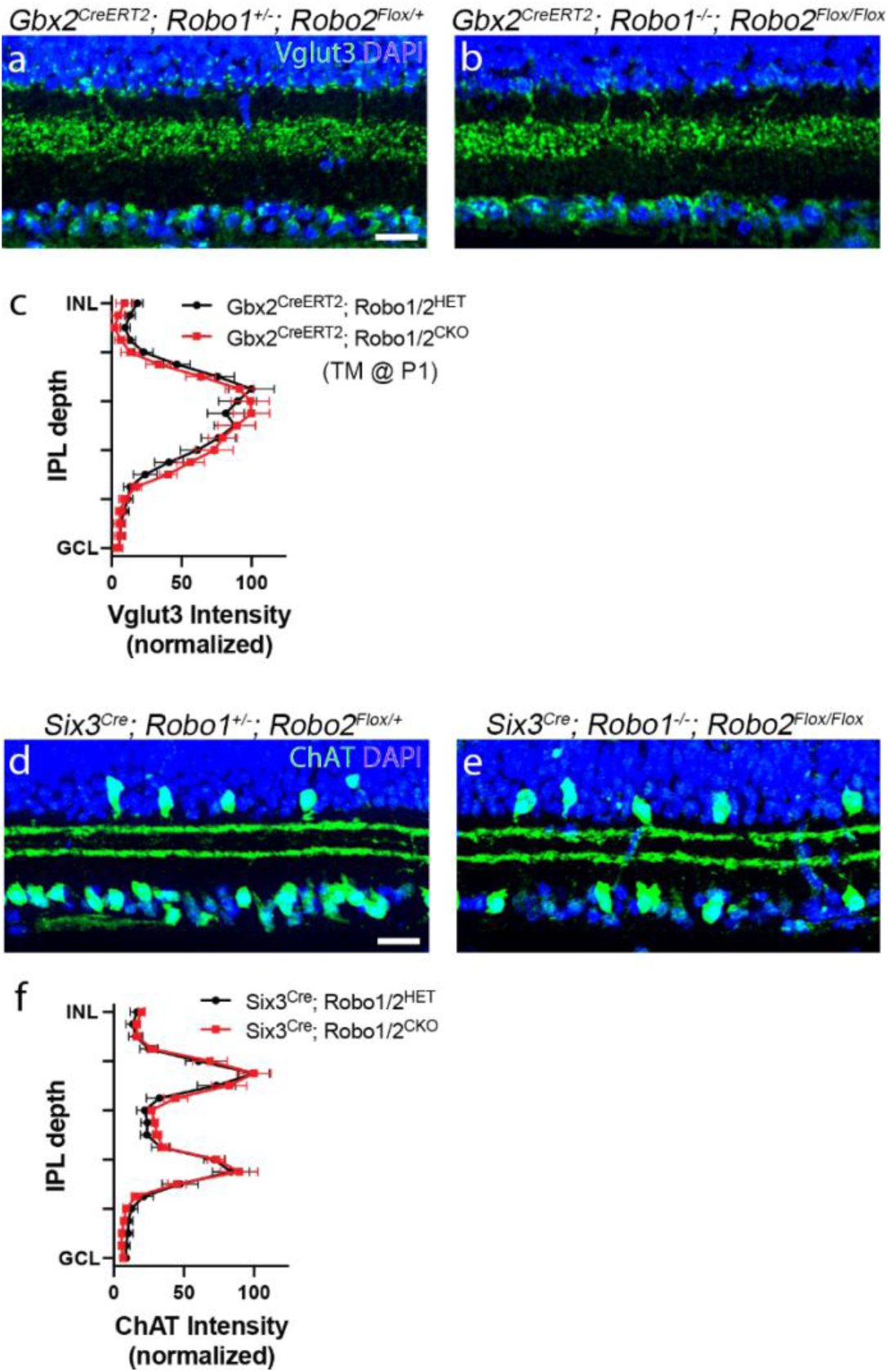
The cell type selective and cell autonomous roles of Robo receptors in retinal amacrine cell development. (**a-c**) In adult retinal sections, the dendrite stratifications of Vglut3+ amacrine cells in heterozygous (**a**) control and (**b**) *Gbx2^CreERT2^; Robo1^-/-^; Robo2^Flox/Flox^* mouse retinas. Tamoxifen (25 µg) was administered at P0 and P1. (**c**) The fluorescent intensity plot of Vglut3 along the IPL depth in control (black) and *Robo1/2 cKO* (red) mice. (**d-f**) The dendrite stratifications of starburst amacrine cells were analyzed by ChAT immunohistochemistry in (**d**) heterozygous controls and (**e**) *Six3^Cre^; Robo1^-/-^; Robo2^Flox/Flox^* conditional knockout mice. (**f**) The fluorescent intensity plot of ChAT along the IPL depth in control (black) and Robo1/2 cKO (red) mice. Scale bar, 50μm in (**a**) and (**d**).

